# Shear Stress Promotes Remodeling of Platelet Glycosylation via Upregulation of Platelet Glycosidase Activity: The Ulterior Cause of MCS-Related Thrombocytopenia?

**DOI:** 10.1101/2024.03.05.583630

**Authors:** Yana Roka-Moiia, Sabrina Lewis, Estevan Cleveland, Joseph E. Italiano, Marvin J. Slepian

## Abstract

**Objective:** Mechanical circulatory support (MCS) is a mainstay of therapy for advanced and end-stage heart failure. Accompanied by systemic anticoagulation, contemporary MCS has become less thrombogenic, with bleeding complications emerging as a major cause of readmission and 1-year mortality of device-supported patients. Shear-mediated platelet dysfunction (SMPD) and thrombocytopenia of undefined etiology are primary drivers of MCS-related bleeding. Recently, it has been demonstrated that deprivation of platelet surface glycosylation is associated with the decline of hemostatic function, microvesiculation, and premature apoptosis. We tested the hypothesis that shear stress induces remodeling of platelet surface glycosylation via upregulation of glycosidase activity, thus facilitating platelet count decline and intense microvesiculation.

**Approach and Results:** Human gel-filtered platelets were exposed to continuous shear stress *in vitro*. Platelets and platelet-derived microparticles were quantified via flow cytometry using size standard fluorescent nanobeads. Platelet surface glycosylation was evaluated using lectin staining and multicolor flow cytometry; lectin blotting was utilized to verify glycosylation of individual glycoproteins. Platelet neuraminidase, galactosidase, hexosaminidase, and mannosidase activities were quantified using 4-methylumbelliferone-based fluorogenic substrates. We demonstrated that shear stress promotes selective remodeling of platelet glycosylation via downregulation of 2,6-sialylation, terminal galactose, and mannose, while 2,3-sialylation remained largely unchanged. Shear-mediated deglycosylation is partially attenuated by neuraminidase inhibitors DANA and zanamivir, strongly suggesting involvement of platelet neuraminidase in observed phenomena. Platelets exhibited high basal hexosaminidase and mannosidase activities; basal activities of platelet neuraminidase and galactosidase were rather low and were significantly upregulated by shear stress. Shear stress of increased magnitude and duration potentiated an incremental decline of platelet count and immense microvesiculation, both being further exacerbated by neuraminidase.

**Conclusions:** Our data indicate that shear stress accumulation, consistent with supraphysiologic conditions of device-supported circulation, promotes remodeling of platelet glycosylation via selective upregulation of platelet glycosidase activity. Shear-mediated platelet deglycosylation is associated with platelet count drop and increased microvesiculation, thus offering a direct link between deglycosylation and thrombocytopenia observed in device-supported patients.

**GRAPHICAL ABSTRACT:** 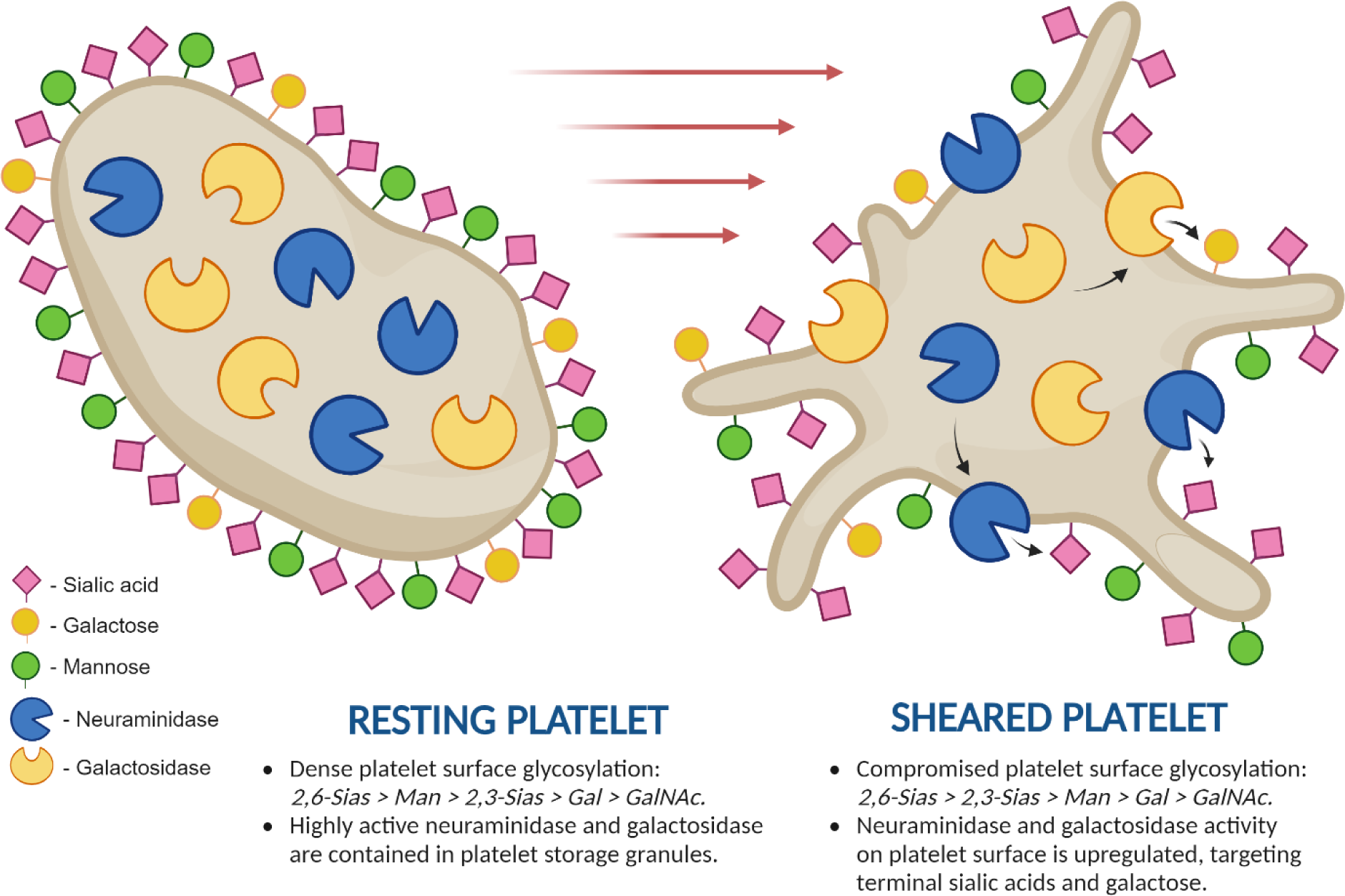

## INTRODUCTION

Mechanical circulatory support (MCS) is a mainstay of therapy for advanced and end-stage heart failure. While effective in restoring hemodynamics, MCS therapy remains limited by serious adverse events related to paradoxical thrombosis and bleeding^1,2^. These adverse events are driven by supraphysiologic shear stress and turbulence imparted to blood cells and proteins, inducing a poorly understood and difficult to manage MCS-related coagulopathy^3,4^. Contemporary MCS accompanied by systemic anticoagulation has become less thrombogenic, putting a major emphasis on bleeding complications^5^. Bleeding is a leading cause of readmission and 1-year mortality for MCS patients: up to 47% of patients on ventricular assist device (VAD) and 65% of patients on extracorporeal membrane oxygenation (ECMO) experience a hemorrhagic event requiring repetitive transfusions and reoperation^6^. The growing body of evidence indicates that shear-mediated quantitative and qualitative platelet dyscrasia is the main driver of MCS bleeding events. Thrombocytopenia defined as platelet count <150,000/μL or 50% less than baseline^5^ is observed in up to 86% of MCS patients^7^. Device-related thrombocytopenia has rapid onset, with platelet count declining drastically within 24 hours following device implantation, and persisting until device removal^8^. The severity of platelet count drop depends on the type of device: veno-arterial ECMO −69.6%, veno-veno ECMO −40.9%, Impella 5.5 −20.9%, and Centrimag biventricular VAD −6.5%^8^. Severely thrombocytopenic MCS patients have increased bleeding risk and longer ICU stays, require frequent platelet transfusions, and are at much higher risk of mortality with relative count drop in day 1 following device initiation being an independent prognostic factor of in-hospital mortality^5,9^.

Platelets are small anucleate blood cells essential for blood coagulation and thrombus formation following vascular damage. When stimulated with biochemical agonists (ADP, thrombin, or collagen), platelets undergo activation, rapid shape change, secretory granule release, and aggregation ultimately forming a thrombus^10^. To date, studies of platelet activation have been largely focused on the role of biochemical stimuli acting via receptor-mediated means. The significance of shear stress as a mechanical force affecting platelet physiology and lifespan has been appreciated rather recently with expanded use of MCS therapy rendering high-speed turbulent flows.^11,12^ Our research group was the first to demonstrate that shear-mediated platelet dysfunction (SMPD) phenotypically differs from platelet activation by biochemical agonists and is characterized by a distinct pattern of orchestrated functional responses^12^. Unlike biochemical agonists, device-related shear stress potentiates platelet pro-apoptotic behavior, such as mitochondrial collapse, membrane depolarization, downregulation of key membrane receptors, abolished aggregatory response to biochemical stimuli, and extensive microvesiculation – all potential contributors to bleeding outcomes^12–16^.

Platelet membrane glycoproteins αIIbβ3 and GPIb-IX are essential for platelet adhesion and aggregation. Binding their protein ligands fibrinogen and vWF, these receptors bridge platelets to the damaged vessel wall and to one another during thrombus formation. Both αIIbβ3 and GPIb-IX are heavily glycosylated, carrying nearly 80% of platelet surface glycosylation^17,18^. Platelets are equipped with enzymatic machinery required to maintain and alter their glycosylation status in an autocrine and paracrine manner^19,20^. Yet, the function of platelet receptor glycans has long been under-appreciated. Over the last decade, evidence has emerged indicating the critical role of glycosylation in regulation of integrin receptor expression and ligand binding^18,21^. Deprivation of platelet surface glycans is associated with a decline of their hemostatic function, microvesiculation, apoptosis, and platelet clearance via phagocytosis by hepatocytes and macrophages^22,23^. Acceleration of platelet deglycosylation during sepsis, viral and microbial infections, or long-term storage in blood bank causes premature platelet clearance, leading to severe thrombocytopenia, bleeding, and lack of hemostatic effect from platelet transfusion^24^. Recent studies indicate that shear force may also affect surface glycosylation of other vascular cells^25^.

Evident similarity of the morphophysiological events triggered by platelet deglycosylation and SMPD led us to hypothesize that device-related shear stress can as well affect platelet glycosylation, thus potentiating platelet dysfunction and device-related bleeding coagulopathy. Here, we test the hypothesis that exposure to elevated shear stress induces remodeling of platelet surface glycosylation via upregulation of platelet glycosidase activity. Specifically, we examine the effect of uniform continuous shear stress of increasing magnitude and duration on platelet surface glycosylation and glycosidase activity. Further, we investigate the origin of platelet glycosidases and identify other potential sources of glycosidase activity in blood. Lastly, we evaluate the cumulative effect of shear stress and exogenous neuraminidase on platelet count and microvesiculation - all potential contributing factors to the pathogenesis of device-related platelet dysfunction and thrombocytopenia.

Overall, we demonstrate that shear stress promotes selective remodeling of platelet glycosylation via downregulation of 2,6-sialylation, terminal galactose and mannose, while 2,3-sialylation remains largely unchanged. Shear-mediated platelet deglycosylation can be partially attenuated by neuraminidase inhibitors 2-deoxy-N-acetylneuraminic acid (DANA) and zanamivir, implicating platelet neuraminidase in observed phenomena. Characterizing platelet glycosidase machinery, we found that platelets exhibit high basal hexosaminidase and mannosidase activities. Notably, basal activities of neuraminidase and galactosidase on platelets were rather low and were significantly upregulated after exposure to shear stress. Lastly, we show that shear stress of increasing magnitude and duration potentiates a substantial drop in platelet count and intense microvesiculation. Neuraminidase further exacerbates shear-mediated decrease of platelet count and microparticle generation, thus offering a direct link between desialylation and thrombocytopenia observed in device-supported patients.

## MATERIALS AND METHODS

The data that support the findings of this study are available from the first author upon reasonable request. For more details of reagents and methods used, please see the major resources tables in the Data Supplement.

### Blood collection and fractionation

The study protocol was approved by the University of Arizona IRB (protocol ID: 1810013264/MOD00001866, from 06/17/2022). Blood was collected from healthy volunteers via venipuncture, anticoagulated with acid citrate dextrose solution and processed to obtain serum, platelet rich-plasma (PRP), and platelet-poor plasma (PPP). Gel-filtered platelets (GFP) were isolated from PRP via gel-chromatography on Sepharose-2B^12^, and platelet count was measured on Z1 Particle Counter (Beckman Coulter, Brea, CA).

### Platelet exposure to shear stress

Recalcified GFP (20,000 or 100,000 plt/µL, 2.5 mM CaCl_2_) was subjected to uniform continuous shear stress in a hemodynamic shearing device, a computer-controlled modified cone-plate-Couette viscometer^26^. Prior to shear exposure, platelets were treated with neuraminidase inhibitors oseltamivir, 2,3- didehydro- DANA and zanamivir, metalloproteinase inhibitor GM6001, or EGTA for 30 min at room temperature with gentle agitation. Sheared platelets were then processed for flow cytometry, lectin blotting, or glycosidase activity.

### Flow cytometric assessment of platelet surface glycosylation, degranulation, and glycoprotein surface expression

To detect terminal sugars on the platelet surface, GFP (20,000 plt/μL, 2.5 mM CaCl_2_) were stained with fluorescein-conjugated lectins: Cy5-*Sambucus Nigra* agglutinin I (SNA) for α-2,6-linked sialic acids (Sias), biotinylated *Maackia Amurensis* lectin II (MAL) and PECy5-streptavidin (0.1 ug/ml) for α-2,3-linked Sias, FITC-*Ricinus Communis* agglutinin I (RCA) and FITC-*Erythrina Cristagalli* agglutinin (ECA) for galactose, FITC-soybean agglutinin (SBA) for *N-* acetylgalactosamine (GalNAc), FITC-*Griffonia Simplicifolia* lectin II (GSL) for *N-* acetylglucosamine (GlcNAc), and FITC-*Lens Culinaris* agglutinin (LCA) for mannose^27^. Alternatively, recalcified GFP were double-stained with anti-CD62P-APC and anti- CD42a-FITC or anti-CD63-PE and anti-CD41-APC, to track platelet α-granule and lysosome release, respectively. Flow cytometry was conducted on a FACSCanto II flow cytometer (BD Biosciences, Franklin Lakes, NJ). Ten thousand events were captured within the stopping gate “Platelets + Microparticles”. Single platelets were distinguished from microparticles based on their forward versus side scatter, as compared with the SPHERO™ Nano Fluorescent Size Standard Kit^13^. Marker-positive platelet and microparticle counts were expressed as % of all events in a joint gate “Platelets + Microparticles”. The arbitrary density of lectin binding and glycoprotein surface expression on platelets and microparticles was calculated as the median fluorescence intensity normalized to the median forward scatter (MFI/MFS)^13,15,28^.

### Lectin blotting of platelet glycoproteins

Platelet lysates were separated by SDS-PAGE in 7.5% gel, transferred to a PVDF membrane, and incubated with biotinylated lectins SNA (1:5000), MAL (1:500), RCA (1:2500), and LCA (1:2500) overnight at 4°C.

To visualize biotinylated lectins, the PVDF membrane was sequentially incubated with peroxidase for 30 min and TMB-blotting solution for 2-5 min until bands develop. Lectin blot images were acquired using a Nikon D3500 DSLR camera (Tokyo, Japan) with an EMART photography lighting kit (Seoul, South Korea) and digitally analyzed using GelAnalyzer 19.1 software (www.gelanalyzer.com).

### Glycosidase activity in blood fractions

Glycosidase activity was measured in PRP, PPP, serum, and recalcified GFP (100,000 plt/μL, 2.5 mM CaCl_2_) before and after shear exposure. For intraplatelet glycosidase activity, aliquots of PRP, non-sheared GFP, and sheared GFP were lysed with RIPA buffer containing proteinase and phosphatase inhibitors at the sample to RIPA buffer ratio of 1:1. Enzymatic activities of neuraminidase, galactosidase, hexosaminidase, and mannosidase were evaluated in acidic and neutral pH using corresponding 4-methylumbelliferone-based fluorogenic substrates. Kinetics of 4-methylumbelliferone release from fluorogenic substrates by glycosidases was recorded as an increase in fluorescence for 2 hours at 37°C using FilterMax™ F5 microplate reader (Ex: 360 nm, Em: 430 nm, Molecular Devices, San Jose, CA). The resulting concentration of 4-methylumbelliferone was determined using a calibration curve, and glycosidase activities were calculated as the initial velocity of 4- methylumbelliferone accumulation (nM per minute)^29^.

### Statistical analysis

Results from 6 to 10 independent experiments with different donors were summarized in figures. All flow cytometry and enzymatic assays samples were run in duplicate and triplicate, respectively. All lectin blots were performed with samples from 7 different donors. The numerical data were tested for normality using the Shapiro-Wilk test, and then statistically analyzed using one-way analysis of variance (ANOVA) followed by Dunnett’s multiple comparisons test and paired t-test for paired comparisons using GraphPad Prism 9 (GraphPad Software). Averages are reported as Mean±SEM. The level of statistical significance is indicated on the figures as *p*<0.05 and *p*<0.01.

## RESULTS

### Shear stress and neuraminidase alters platelet surface glycosylation

To test whether shear stress and neuraminidase affect platelet surface glycosylation, we analyzed the surface distribution of terminals sugars, such as Sias, galactose, mannose, GalNAc, and GlcNAc, on platelets using fluorescein-conjugated lectins and flow cytometry. We found that resting platelets exhibited very high basal levels of 2,6- sialylation, as indicated by the high-density binding of SNA lectin (**Fig. 1A**); 2,3- sialylation was less pronounced, as MAL binding density was nearly 4-fold lower as compared to SNA (**Fig. 1D**). Exposure to increasing shear stress resulted in a progressive decrease of platelet 2,6-sialylation, as indicated by the significant decrease of SNA binding on platelets subjected to 70 dyne/cm^2^ shear for 10 min: 3.28±0.28 AU vs. 4.49±0.29 AU on resting platelets (p≤0.01). The MAL binding detecting 2,3- sialylation remained unchanged after exposure to shear stress. Neuraminidase further exacerbated the decrease of platelet 2,6-sialylation independent of shear exposure, while 2,3-sialylation was not significantly altered (**Fig. 1B, 1E**). Interestingly, the levels of SNA and MAL binding on PDMPs generated by shear were on average 3.5-fold higher than on platelets, suggesting the increased density of 2,6- & 2,3-sialylation on PDMPs (**Fig. 1C<u>, 1F</u>**).

**Figure 1.**
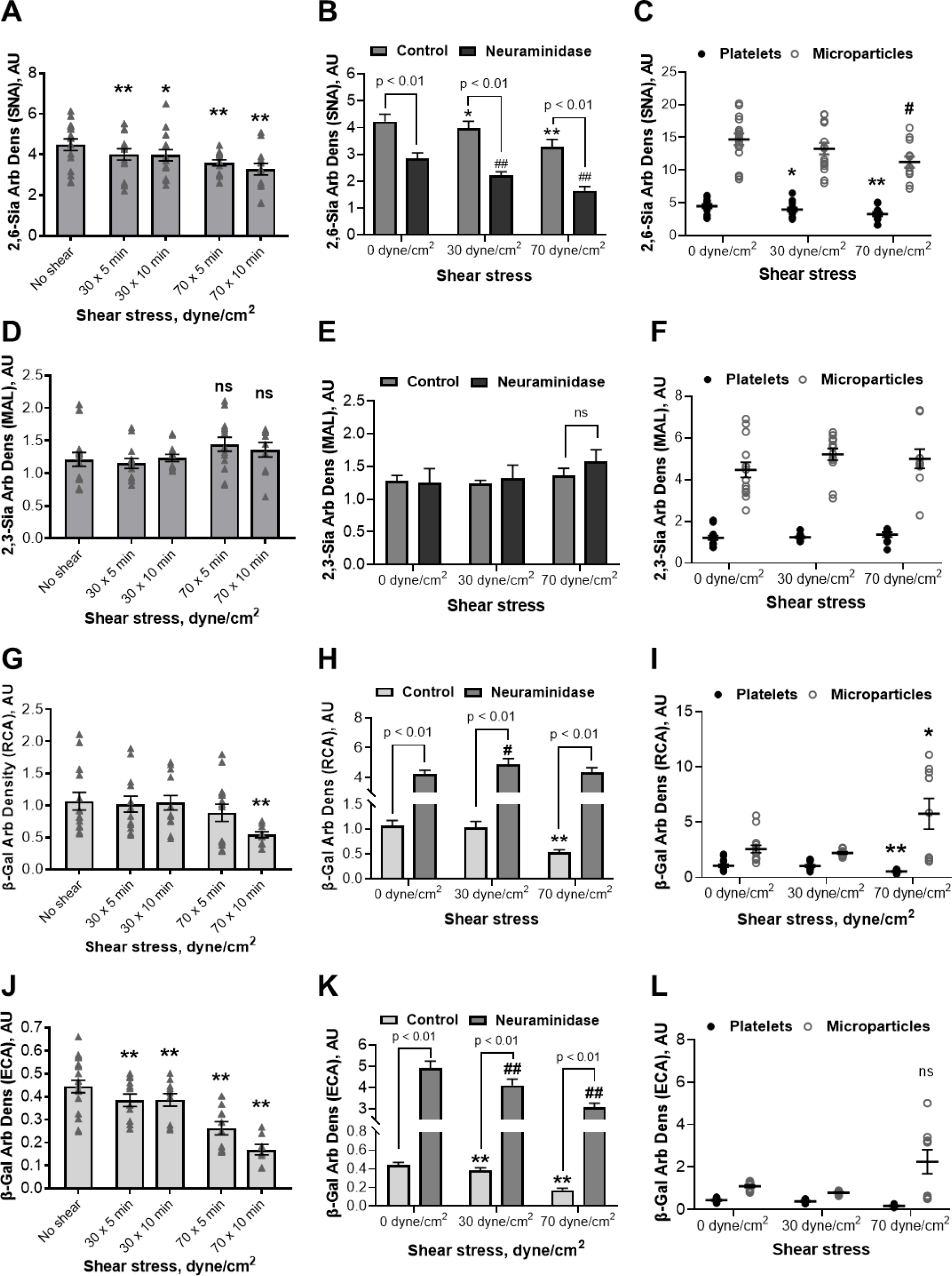
Shear stress promotes a decrease of 2,6-sialylation and galactosylation on platelets, while 2,3-sialylation remains unaltered. Enzymatic desialylation by neuraminidase is associated with the downregulation of 2,6-sialylation and an increase of platelet surface galactose. A, B – arbitrary density of 2,6-sialic acids on platelets (n = 8), D, E – arbitrary density of 2,3-sialic acids on platelets (n = 8), C, F – arbitrary density of 2,6- and 2,3-sialic acids on platelets and microparticles (n = 8); G, H – arbitrary density of galactose on platelets detected using RCA lectin (n = 7), J, K – arbitrary density of galactose on platelets detected using ECA lectin (n = 6), I, L – arbitrary density of galactose on platelets and microparticles detected using RCA & ECA lectins (n = 6-7). Mean±SEM, 1-way ANOVA followed by Dunnett multiple comparisons test: *, # - p<0.05, **, ## - p<0.01 vs correspondent unsheared group.

Next, we demonstrated that resting platelets presented galactose-capped glycans on their surface as indicated by the medium-intensity binding of RCA and ECA lectins (**Fig. 1G<u>, 1J</u>**). Exposure to shear stress resulted in an incremental decrease of RCA and ECA binding, indicating the decrease of surface galactose with the increase of shear duration and intensity. Neuraminidase however induced a striking increase of surface galactose on platelets independent of shear exposure (**Fig. 1H<u>, 1K</u>**). When exposed to shear, neuraminidase-treated platelets exhibited decreased levels of ECA and RCA binding. Interestingly, the density of galactose-capped glycans on PDMPs generated by low shear stress was comparable to that on platelets, while 70 dyne/cm^2^ sheared PDMPs exhibited significantly higher RCA & ECA binding not characteristic of those on resting and sheared platelets (**Fig. 1I**, **<u>1L</u>**).

Evaluating GalNAc and GlcNAc, we found that the resting platelets expressed very low levels of these sugars, as indicated by the low-density binding of SBA and GSL lectins (**Fig. 2A**, **<u>2D</u>**). Shear stress did not significantly affect platelet *GalNAc*, while *GlcNAc* density increased 1.8-fold after 70 dyne/cm^2^ shear. Neuraminidase alone rendered a 56-fold increase in *GalNAc* density on platelets (**Fig. 2B**). Yet, when neuraminidase-treated platelets were exposed to shear stress, the levels of *GalNAc* significantly decreased. Neuraminidase also increased *GlcNAc* levels on platelets, but exposure of neuraminidase-treated platelets to shear had no significant effect on GSL binding (**Fig. 2E**). The *GalNAc* and *GlcNAc* density on sheared PDMPs were significantly than on platelets (**Fig. 2C**<u>, **2F**</u>).

**Figure 2.**
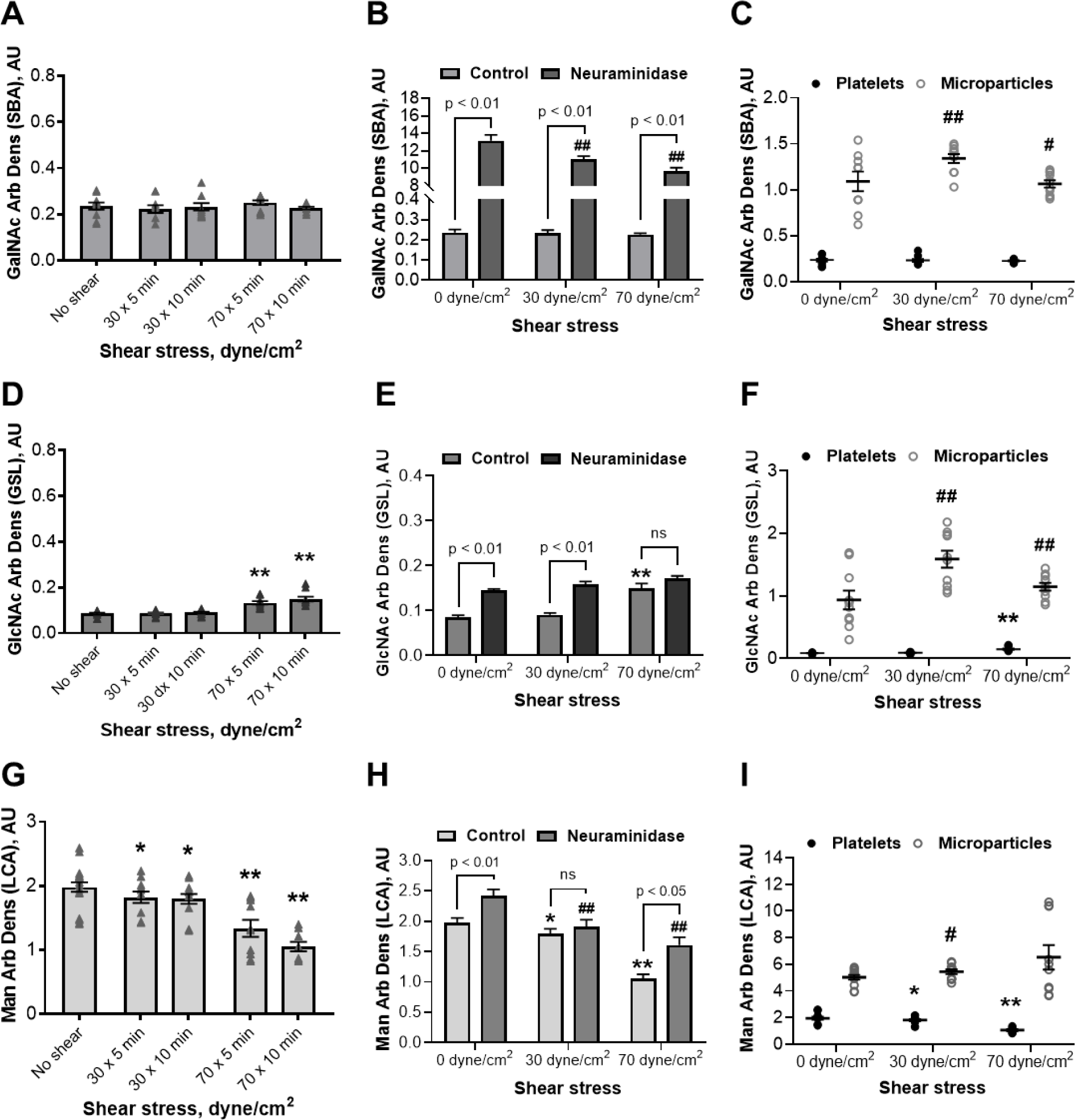
Shear stress promotes a decrease of platelet surface mannose, while N-acetylgalactosamine and N-acetylglucosamine remain largely unaltered. Enzymatic desialylation by neuraminidase is associated with an increase of platelet surface N-acetylgalactosamine and, to a lesser extent, N-acetylglucosamine and mannose. A, B – arbitrary density of N-acetylgalactosamine on platelets (n = 8), D, E – arbitrary density of N-acetylglucosamine on platelets (n = 8), G, H – arbitrary density of mannose on platelets (n = 6), C, F, and I – arbitrary density of N-acetylgalactosamine, N-acetylglucosamine, and mannose on platelets and microparticles (n = 8). Mean±SEM, 1-way ANOVA followed by Dunnett multiple comparisons test: *, # - p<0.05, **, ## - p<0.01 vs correspondent unsheared group.

Lastly, we showed that resting platelets exhibited high levels of mannose-capped glycans, as indicated by high-density binding of LCA lectin (**Fig. 2G**). Exposure to shear stress resulted in a significant decrease of platelet surface mannose, as shown by the progressive decrease of LCA binding with increasing shear magnitude and duration. Neuraminidase treatment resulted in a modest, yet statistically significant increase of platelet surface mannose (**Fig. 2H**). Exposure of neuraminidase-treated platelets to shear rendered a 1.5-fold decrease in platelet LCA binding. The levels of LCA binding on PDMPs were 3-6-fold higher than on platelets in corresponding groups, suggesting increased density of mannose on PDMPs (**Fig. 2I**).

### Shear stress & neuraminidase promote deglycosylation of platelet glycoproteins

To evaluate the effect of shear stress and neuraminidase on glycosylation levels of major platelet glycoproteins, we blotted whole-platelet lysates with four lectins showing the highest binding intensity on platelets as indicated by flow cytometry: SNA for 2,6- linked Sias, MAL for 2,3-linked Sias, RCA for galactose, and LCA for mannose. We found that numerous platelet glycoproteins are extensively 2,6-sialylated, as indicated by a dozen of distinct protein bands (50-400 kDa) revealed by SNA in resting platelet lysates (**Fig. 3A**). Exposure to shear stress resulted in a drastic decrease in 2,6-sialylation, as seen from severe fading of all protein bands. Similarly, neuraminidase-mediated desialylation resulted in the dissipation of most SNA-stained bands, yet several distinct protein bands appeared not characteristic of resting platelets. Neuraminidase inhibitors DANA and zanamivir (but not oseltamivir) as well as EGTA attenuated shear-mediated decrease of platelet 2,6-sialylation as shown by restoration of the most if not all SNA-stained bands. Pan-metalloproteinase inhibitor GM6001 only partially restored glycoprotein sialylation post-shear.

**Figure 3.**
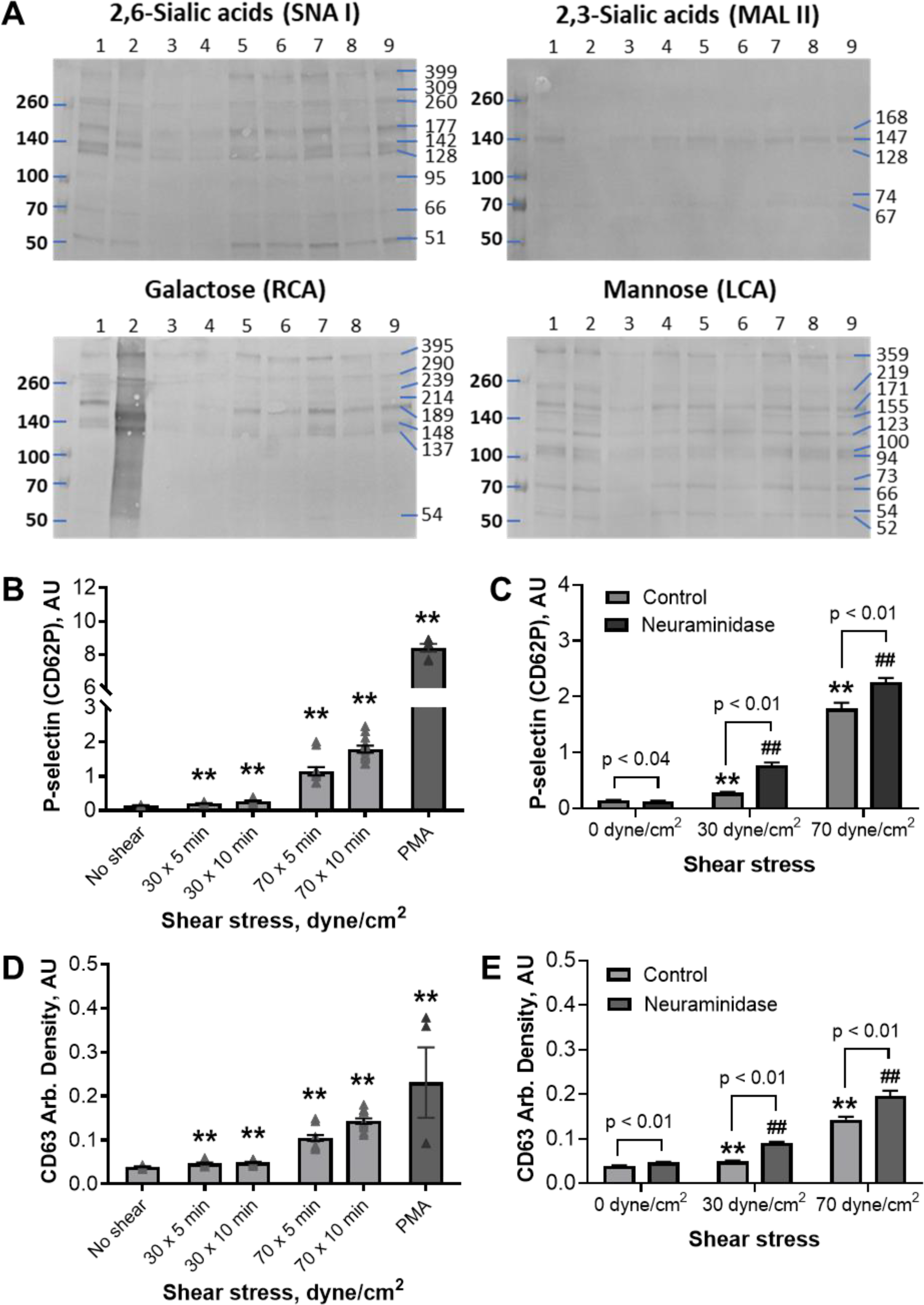
Shear stress and neuraminidase promote deglycosylation of platelet glycoproteins, associated platelet degranulation, and lysosomal release: A – representative lectin blotting of platelets exposed to shear stress and neuraminidase; B, C – arbitrary density of P-selectin on platelets exposed to shear stress and neuraminidase (n = 6); D, E - arbitrary density of CD63 on platelets exposed to shear stress and neuraminidase (n = 6). Protein load for lectin blots: SNA I and RCA - 6 μg/well, MAL II and LCA – 7 μg/well. Well order: 1 and 9 – intact platelets, 2 – 0.1 U/mL neuraminidase, 3 – 70 dyne/cm^2^ shear stress for 30 min, 4 – 1 mM oseltamivir + shear stress, 5 – 1 mM DANA + shear stress, 6 – 1 mM zanamivir + shear stress, 7 – 2 mM EGTA + shear stress, 8 – 100 nM GM6001 + shear stress. Mean±SEM, 1-way ANOVA followed by Dunnett multiple comparisons test: **, ## - p<0.01 vs correspondent unsheared group.

Analyzing 2,3-sialylation of platelet glycans, we found that in resting platelets only 4-5 protein bands were revealed by MAL (**Fig. 3A**), suggesting that, unlike 2,6-sialylation, 2,3-sialylation is rather unique motif within selected platelet glycoproteins. Following shear stress and neuraminidase, only one faint protein band could be detected (152 or 69 kDa, respectively). Neuraminidase inhibitors, EGTA, and GM6001 largely recovered 2,3-sialylation of high-molecular-weight bands and to a lesser extent low-molecular-weight bands post-shear.

The RCA blotting revealed an abundance of galactose-capped high-molecular-weight platelet glycoproteins (**Fig. 3A**). Neuraminidase induced massive desialylation as indicated by the increased density of RCA-stained bands and their redistribution towards lower molecular weights. Shear stress, however, resulted in substantial fading of most protein bands and the decrease of molecular weight for several others. Neuraminidase inhibitors DANA and zanamivir (but not oseltamivir), GM6001, and EGTA partially restored 4-8 protein bands out of 8 bands observed in resting platelets.

Lastly, the LCA blotting showed that mannose-capped glycans were common in platelet glycoproteins of various molecular weights (**Fig. 3A**). At least 12 distinct protein bands were revealed by LCA in resting platelets. Neuraminidase-mediated desialylation did not cause any apparent decrease in mannose levels, instead several protein bands showed increased density as compared to those in control (66 and 120 kDa). Shear stress resulted in extensive fading or complete disappearance of LSA-stained bands. Neuraminidase inhibitors rendered partial recovery of 9-11 protein bands post-shear, with DANA showing the highest efficacy. Metalloproteinase inhibitor GM6001 and, to a greater extent, EGTA resulted in recovery of all 12 glycoprotein bands, but their density remained compromised.

### Shear stress and neuraminidase facilitate mild platelet degranulation and lysosomal release

To test whether shear stress and neuraminidase promote platelet degranulation, we monitored exposure of corresponding marker proteins on platelet surface: P selectin – for α-granule and CD63 – for lysosomal release. We found that shear stress promoted the progressive increase of P-selectin density on platelets as the shear stress duration and magnitude increased (**Fig. 3B**). Neuraminidase alone did not affect platelet surface P-selectin (**Fig. 3C**). Yet, when neuraminidase-treated platelets were exposed to shear stress, they exhibited substantially higher surface P-selectin levels as compared to untreated counterparts: 0.28±0.02 vs 0.77±0.05 AU and 1.79±0.10 vs 2.26±0.07 AU, for 30 and 70 dyne/cm^2^ shear, respectively. We also demonstrated that shear stress did not substantially alter platelet CD63 density (**Fig. 3D**). Similarly, neuraminidase alone did not induce evident alteration of platelet CD63 (**Fig. 3E**). Yet, neuraminidase-treated platelets demonstrated significantly higher surface levels of CD63 following exposure to 30 and 70 dyne/cm^2^ shear stress than correspondent control groups.

### Glycosidase activity in platelets, blood plasma and serum

To test whether shear stress promotes upregulation of platelet glycosidase activity, we measured enzymatic activities of neuraminidase, galactosidase, hexosaminidase, and mannosidase. Activities of platelet extracellular glycosidases (intact GFP) and intracellular glycosidases (lysed GFP) were measured in neutral and acidic pH to differentiate between two pH-sensitive enzyme fractions potentially involved in shear-mediated deglycosylation.

We found that basal activity of extracellular neuraminidase on platelets was very low for both neutral and acidic fractions. Platelet exposure to shear stress resulted in an 8-fold increase of *neutral* extracellular neuraminidase activity, while *acidic* extracellular neuraminidase remained unaltered (**Fig. 4A**). Basal activity of *neutral* intracellular neuraminidase was even higher than extracellular neuraminidase and further increased dramatically post-shear: 0.24±0.02 vs 0.73±0.12 nM/min in unsheared and sheared lysed GFP, respectively. *Acidic* intracellular neuraminidase activity was also higher than its extracellular levels, but unlike *neutral* neuraminidase, it did not increase after shear (**Fig. 4A**). Analyzing galactosidase activity, we found that resting platelets exhibited detectable levels of both neutral and acidic extracellular galactosidase activities, and exposure to shear stress resulted in a moderate increase of these activities (**Fig. 4B**).

**Figure 4.**
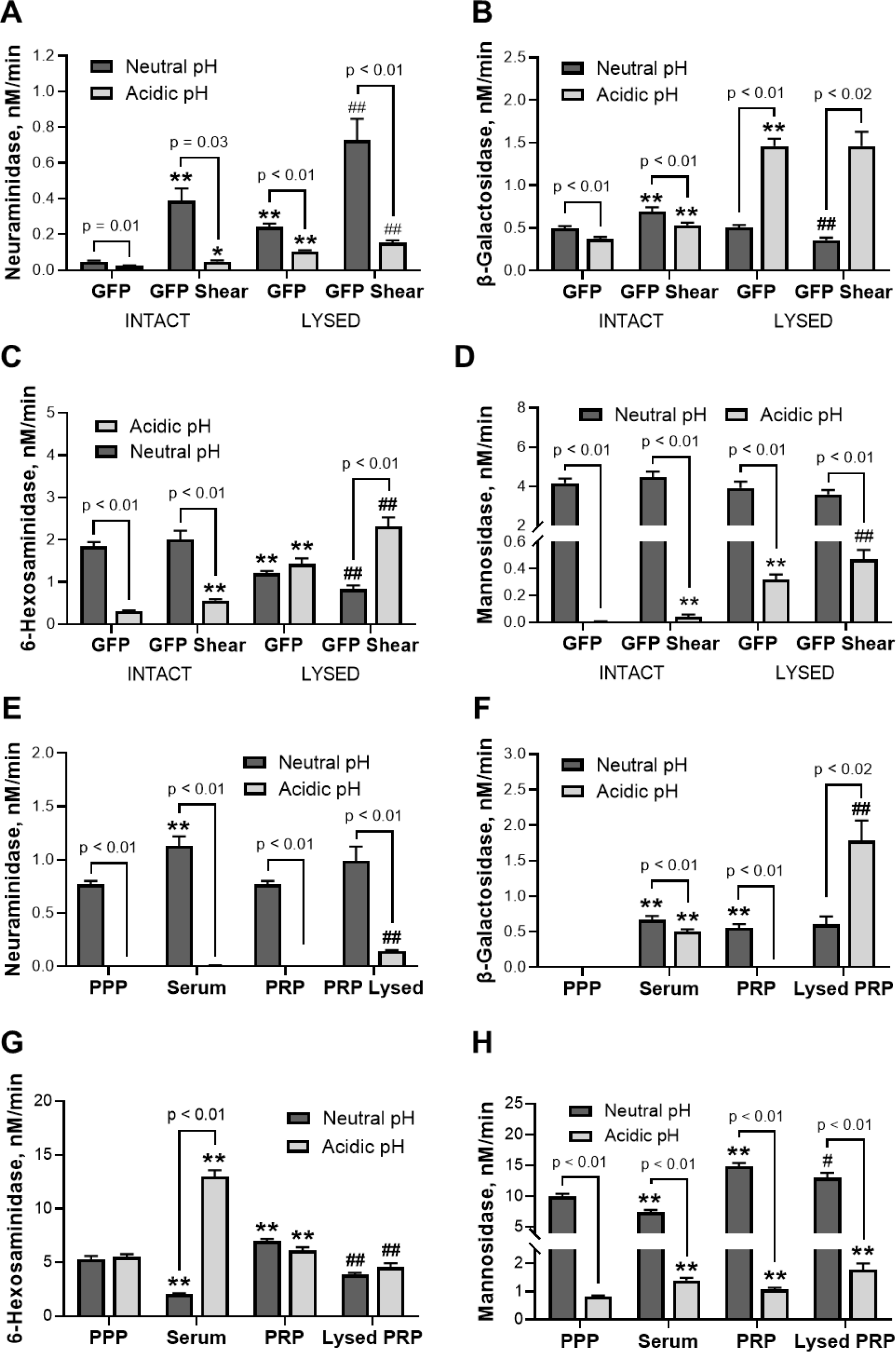
Neutral and acidic glycosidase activity in platelets, blood plasma, and serum. A, B, C, and D – enzymatic activities of neuraminidase (n = 8), galactosidase (n = 8), hexosaminidase (n = 6-8), and mannosidase (n = 7-9) in platelets before and after shear stress; E, F, G and H – enzymatic activity of neuraminidase (n = 10), galactosidase (n = 10), hexosaminidase (n = 11), and mannosidase (n = 14-15) in plasma, serum and platelet-rich plasma. Mean±SEM, 1-way ANOVA followed by Dunnett multiple comparisons test: *, # - p<0.05, **, ## - p<0.01 vs correspondent unsheared group.

Basal activity of *acidic* intracellular galactosidase was 4-fold higher than its extracellular levels (1.46±0.09 nM/min in lysed GFP vs 0.37±0.03 nM/min in intact GFP) and remained elevated post-shear.

We also showed that resting platelets exhibited very high basal activity of *neutral* extracellular hexosaminidase, which remained unaltered post-shear (**Fig. 4C**). In contrast, basal activity of *acidic* extracellular hexosaminidase was rather low, increasing 1.8 times following shear exposure. In lysed GFP, *acidic* intracellular hexosaminidase activity was significantly higher than its extracellular levels (1.42±0.13 vs 0.31±0.01 nM/min), further increasing 1.6 times post-shear. Evaluating mannosidase activity, we showed that all intact and lysed platelet groups exhibited very high *neutral* extracellular mannosidase activity (**Fig. 4D**). Yet virtually no *acidic* extracellular mannosidase activity was detected in intact GFP before or after shear exposure. Basal activity of *acidic* intracellular mannosidase in lysed GFP was rather low and slightly increased post-shear.

Next, glycosidase activity was accessed in blood plasma, serum, and PRP (intact and lysed) to identify alternative sources of glycosidases that might affect platelet glycosylation in the bloodstream. We identified substantial levels of *neutral*, but not acidic, neuraminidase activity in PPP and PRP (**Fig. 4E**). Traceable *acidic* neuraminidase activity was only detected in lysed PRP likely of platelet origin. The *neutral* neuraminidase activity in lysed PRP increased as compared to intact PRP. A dramatic increase of *neutral* neuraminidase activity was also detected in serum (1.13±0.09 nM/min in serum vs 0.77±0.03 in PPP), likely associated with neuraminidase release from platelets during clotting. No measurable galactosidase activity was detected in plasma, neither in neutral nor in acidic pH (**Fig. 4F**). Yet, *neutral* galactosidase activity was detected in PRP at levels comparable to intact and lysed GFP. Very high levels of *acidic* galactosidase activity were detected in lysed PRP (1.78±0.29 nM/min), and not in intact PRP. Unlike plasma, serum showed tracible galactosidase activity likely originating from platelets and other blood cells.

Hexosaminidase activity in plasma was much higher than galactosidase and neuraminidase altogether (**Fig. 4G**); both neutral and acidic hexosaminidase activities were detected. PRP exhibited even higher hexosaminidase activity, likely of platelet origin. Interestingly, serum showed comparably low activity of *neutral* hexosaminidase, while activity of *acidic* fraction was very high, nearly 3-fold higher than in PPP. *Neutral* mannosidase activity across all plasma fractions was very high, greatly exceeding even hexosaminidase levels (**Fig. 4H**). As expected, PRP demonstrated the highest neutral mannosidase activity due to an additive effect associated with the platelet pool of this enzyme. However, enzymatic activity of *acidic* mannosidase in plasma was much lower, with the highest activity detected in lysed PRP. Serum showed significantly lower *neutral* mannosidase activity than plasma, nevertheless *acidic* mannosidase activity was nearly twice as high as in PPP.

### Shear stress & neuraminidase induce platelet count drop and generation of platelet-derived microparticles

To evaluate the effect of shear stress and neuraminidase on platelet count and microvesiculation, platelets and PDMPs were identified using two platelet-specific markers CD41 and CD42a and quantified by flow cytometry. We demonstrated that shear stress of increased magnitude and duration resulted in a progressive decrease of platelet count (**Fig. 5A**, **<u>5G</u>**). Neuraminidase-treated platelets demonstrated even higher sensitivity to shear stress, with CD41+ and CD42a+ platelet counts decreasing more drastically as compared to untreated control groups exposed to shear stress of the same magnitudes and durations (**Fig. 5B**, **<u>5H</u>**).

**Figure 5.**
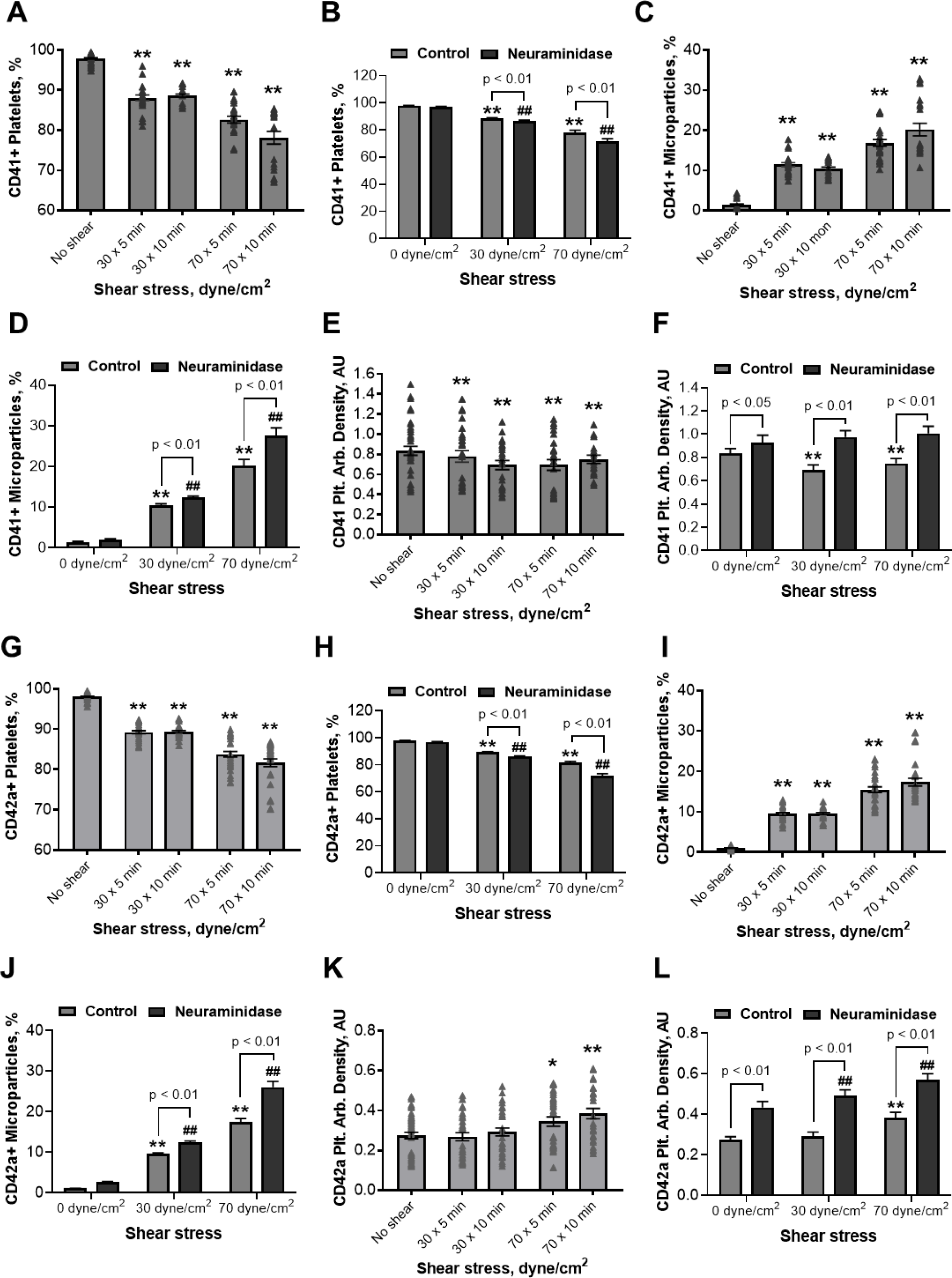
Increasing shear stress and neuraminidase promote a progressive decrease of platelet count and intense microvesiculation. Shear stress rendered a modest decrease of α2bβ3 arbitrary density and an increase in GPIX. Neuraminidase however upregulated the surface expression of both receptors. A, B – the number of CD41+ platelets, C, D – the number of CD41+ microparticles, E, F – α2bβ3 arbitrary density on platelets; G, H - the number of CD42a+ platelets, I, J – the number of CD42a+ microparticles, K, L - GPIX arbitrary density on platelets. n = 8, Mean±SEM, 1-way ANOVA followed by Dunnett multiple comparisons test: * - p<0.05, **, ## - p<0.01 vs correspondent unsheared group.

We also showed that shear stress promotes generation of PDMPs carrying platelet specific markers CD41 and CD42a (**Fig. 5C** & **<u>5I</u>**). As such, platelet exposure to 30 dyne/cm^2^ shear stress for 10 min rendered 10.5±0.4% of CD41+ and 9.6±0.2% of CD42+ PDMPs, while 70 dyne/cm^2^ shear rendered 20.2±1.6% CD41+ and 17.4±1.0% CD42a+ PDMPs. Pre-treatment with exogenous neuraminidase resulted in increased generation of PDMPs following platelet exposure to shear stress, more evident when higher shear magnitudes were applied (**Fig. 5D** & **<u>5J</u>**).

Lastly, we evaluated the surface density of CD41 and CD42a on platelets following exposure to increased shear stress and neuraminidase. We demonstrated that shear stress induced a decrease of CD41 density on platelets with increasing shear magnitude and duration (**Fig. 5E**). Neuraminidase induced an evident increase of CD41 surface density in all platelet groups, independently of shear exposure (**Fig. 5F**). As such, the percent increase for neuraminidase treated platelets as compared to untreated platelets was 12.0%, 42.0%, and 33.3%, for unsheared, 30 and 70 dyne/cm^2^ sheared groups, respectively. Analyzing CD42a surface density, we found that 70 dyne/cm^2^ shear stress induced a mild increase of platelet CD42a (**Fig. 5K**).

Interestingly, neuraminidase alone also induced a substantial increase of platelet CD42a surface density in unsheared and sheared platelets alike (**Fig. 5L**). As such, a neuraminidase-mediated upregulation of platelet CD42a surface expression was 59.2%, 69.0%, and 50% in unsheared, 30 and 70 dyne/cm^2^ sheared groups, respectively, as compared to untreated controls.

## DISCUSSION

In our attempt to identify triggering mechanisms of SMPD and MCS-related thrombocytopenia, we study the effect of shear stress on platelet glycosylation and glycosidase activity, evaluate alternative sources of glycosidases in blood plasma, and test the cumulative effect of platelet exposure to shear stress and exogenous neuraminidase on platelet count and microvesiculation. We report that shear stress of increasing magnitude and duration potentiates selective remodeling of platelet surface glycosylation via downregulation of 2,6-sialylation associated with the decrease in terminal galactose and mannose; simultaneously, 2,3-sialylation, GalNAc, and GlcNAc levels on platelets remain unaffected by shear. In comparison, enzymatic desialylation induced by neuraminidase renders a similar decrease of 2,6-sialylation accompanied by the striking increase in terminal galactose, GalNAc, and to a lesser extent GlcNAc and mannose on platelets. Shear-mediated deglycosylation of platelet glycoproteins can be partially attenuated by neuraminidase inhibitors DANA and zanamivir, implicating platelet neuraminidase in observed phenomena. Indeed, we demonstrate that platelets possess highly active extracellular and intracellular glycosidases. Some platelet glycosidases are constitutively active (hexosaminidase and mannosidase), while others are upregulated by shear stress (neuraminidase and galactosidase) – all potential contributors to shear-mediated deglycosylation. All four extracellular platelet glycosidases demonstrate higher enzymatic activity in neutral pH. Among intracellular glycosidases, galactosidase, hexosaminidase, and mannosidase are active in acidic pH suggesting their primarily lysosomal localization, while neuraminidase still favors neutral pH likely residing in alternative storage granules. Beyond platelets, blood plasma offers a rich source of active neuraminidase, hexosaminidase, and mannosidase. Lastly, we show that shear stress of increasing magnitude and duration potentiates a substantial platelet count decline and intense microvesiculation. Neuraminidase further exacerbates the shear-mediated decrease of platelet count and intensifies microparticle generation, thus offering a direct link between platelet desialylation and thrombocytopenia observed within device-supported circulation.

Platelet surface glycosylation or “sugar coat” is composed of glycosaminoglycans associated with glycoprotein receptors abundantly expressed on the platelet membrane. Glycosaminoglycan moieties are covalently linked to their proteins via N-glycosidic bond (N-glycans) or O-glycosidic bond (O-glycans). *N-glycans* begin from N-acetylglucosamine (GlcNAc) β-linked to asparagine; GlcNAc is further extended with sugar branches made of galactose, poly-mannose, N-acetyllactosamine or fucose (**Fig. 6A**). N-glycans could be capped by Sias, but also galactose or fucose^24^. *O-glycans* begin with N-acetylagalactosamine (GalNAc) α-linked to serine or threonine; GalNAc is then extended with galactose, GlcNAc, fucose, or Sias^24^ (**Fig. 6A**). Sia caps are attached to terminal glycan structures via α2,3-, α2,6- or α2,8-linkages. Our findings indicate that surface glycosylation of resting platelets is composed of 2,6-sialylated and to a lesser extent 2,3-sialylated glycans. Curiously, mannose- and galactose-capped glycans are also abundantly present on resting platelets, yet virtually no surface GlcNAc or GalNAc were detected. The glycosylation pattern of resting platelet sugar coat could be summarized as *2,6-Sias* > Man > *2,3-Sias* > Gal > GalNaAc > GlcNAc. Lectin blotting of resting platelet lysates revealed similar glycan distribution trends. The majority of platelet glycoproteins were extensively sialylated, with 2,6-Sia-capped glycans significantly outnumbering 2,3-Sia-capped glycans. Many glycans lacked terminal sialylation and were capped by terminal mannose or galactose instead. Comparing observed molecular weights of lectin-stained protein bands with molecular weights, expression^30^ and glycosylation levels^24^ of major platelet glycoprotein receptors, we can speculate on the primary contributors to platelet surface glycosylation (see Supplemental Materials, **Table I**).

**Figure 6.**
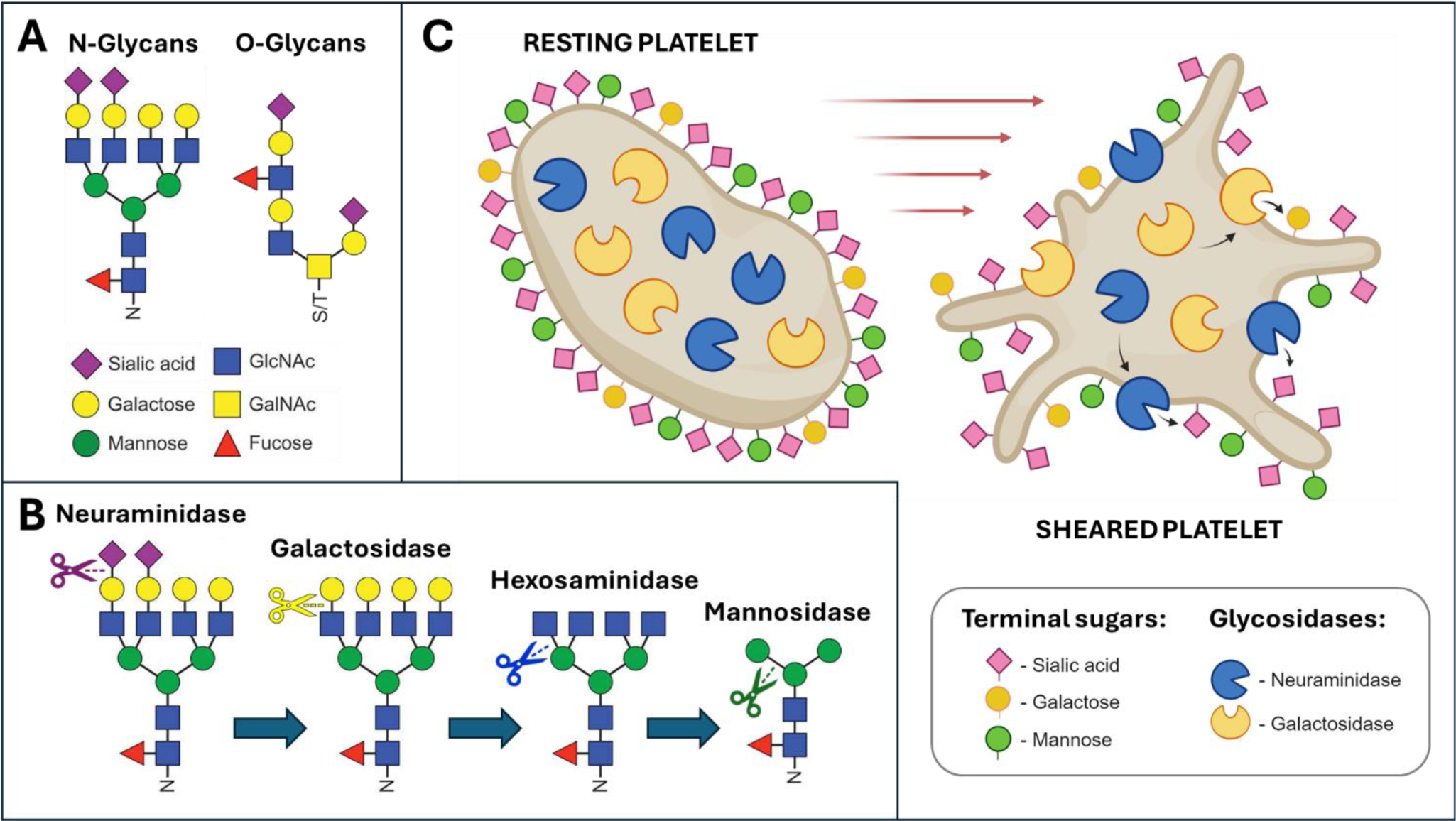
Shear stress potentiates platelet surface deglycosylation via upregulation of glycosidase activity on platelets: A – composition of platelet N- and O-linked glycans; B – sequential deglycosylation of terminal sugars in platelet glycans by neuraminidase, galactosidase, hexosaminidase, and mannosidase; C – potential mechanism of shear-mediated upregulation of neuraminidase and galactosidase activity on platelets and subsequent remodeling of platelet surface glycosylation by the platelet glycosidases.

Sias and other capping sugars are attached to both N- and O-glycans via α- linkage, so they are conformationally distant from the core sugar branches and presented for ligand binding and enzymatic cleavage^24^. Enzymatic desialylation of platelet glycans results in uncapping of underlying sugar moieties, so they become accessible and could potentially be removed by correspondent glycosidases. Comparing the effect of enzymatic desialylation by neuraminidase and shear stress on platelet surface glycosylation, we demonstrated that neuraminidase induced a notable decrease of 2,6-sialylation on platelets surface, associated with the striking increase of terminal galactose and GalNAc levels likely caused by uncapping of these underlying sugars previously masked by Sias. As such, the sugar coat pattern of neuraminidase-treated platelets significantly differs from resting platelets: GalNAc > Gal > *2,6-Sias* > Man > *2,3-Sias* > GlcNAc. Lectin blotting of neuraminidase-treated platelets confirmed the decrease in 2,6- and 2,3-desialylation of most, if not all, platelet glycoproteins accompanied by presentation of galactose- and mannose-capped glycans and redistribution of protein bands towards lower molecular weights – all indicators of deglycosylation. It has been previously reported that bacterial and viral sialidases promote platelet desialylation and an associated increase in terminal galactose and GlcNAc^31,32^.

Next, we found that, similarly to neuraminidase treatment, exposure to increased shear stress rendered a substantial decrease in platelet 2,6-sialylation, and the levels of galactose- and mannose-capped glycans on platelets also decreased significantly. When neuraminidase-treated platelets were exposed to shear stress, platelet 2,6- sialylation plummeted even further, indicating that shear stress and exogenous neuraminidase impose a cumulative (or synergistic) negative effect on platelet surface glycosylation, potentially implicating neuraminidase in shear-mediated desialylation. Lectin blotting of sheared platelets confirmed aggressive desialylation of all observed glycoproteins. Galactose- and mannose-capped glycan bands were also significanly diminished, confirming our flow cytometry observations. Overall, the glycan surface density on sheared platelets was substantially decreased resulting in the following sugar coat pattern: *2,6-Sias* > *2,3-Sias* > Man > Gal > GalNaAc > GlcNAc. Taken together our observations indicate that shear-mediated platelet desialylation, unlike one mediated by neuraminidase, is associated with a decrease of galactose- and mannose-capped glycans in resting platelets and galactose-, mannose- and GalNAc-capped glycans in neuraminidase-treated platelets following shear exposure. Platelet desialylation by endogenous neuraminidases has been demonstrated in immune thrombocytopenia, Bernard-Soulier syndrome, coronary heart disease, and in patients with acute myocardial infarction.^33,34^ We speculate that shear stress could also upregulate the activity of platelet neuraminidases and facilitate desialylation of platelet glycoproteins, triggering a “domino effect” in the form of a deglycosylation cascade of underlying glycans as sequential removal of terminal galactose, GalNAc, and mannose from platelet glycoproteins by correspondent glycosidases in platelet milieu (**Fig. 6B**). On the other hand, we have previously demonstrated that increased shear stress facilitates downregulation and shedding of several platelet receptors, including GPVI, PECAM-1, and P-selectin,^13,15^ potentially contributing to deglycosylation of platelet surface glycoproteins via proteolytic cleavage of their glycosylated ectodomains.

To assess contributions of neuraminidase-mediated desialylation and metalloproteinase-mediated proteolysis to deglycosylation of platelet glycoproteins under shear, platelets were pretreated with neuraminidase inhibitors, metalloproteinase inhibitor GM6001 or calcium chelator EGTA prior to shear exposure. The lectin blotting of sheared platelets indicated that neuraminidase inhibitors DANA and zanamivir (but not oseltamivir) largely restored 2,6- and 2,3-sialylation of platelet glycoproteins compromised by shear stress to near pre-shear levels, thus verifying the critical role of neuraminidase activity in shear-mediated platelet desialylation. Curiously, the restoration of platelet glycan sialylation by neuraminidase inhibitors coincided with recovery of galactose- and mannose-capped glycan bands in sheared platelets. Likewise, both inhibitors of glycoprotein shedding GM6001 and EGTA partially recover 2,6- and 2,3-sialylation of platelet glycoproteins. The high efficacy of neuraminidase inhibitors in restoring platelet glycan sialylation after shear confirms that shear-mediated desialylation of platelet glycoproteins could be attributed to upregulation of platelet neuraminidase activity. The partial recovery of protein glycosylation by the metalloproteinase inhibitor also suggests that shear-mediated enzymatic shedding of glycosylated ectodomains is in part to blame for the dramatic decrease of platelet glycosylation by shear. In the murine model, it was previously demonstrated that GM6001 effectively mitigated degalactosylation of refrigerated platelets caused by desialylation and subsequent ADAM17-mediated proteolytic shedding of GPIb ectodomain.^35^ Taking into consideration that neuraminidase inhibitors were able to recover glycosylation of Sia-capped glycans as well as galactose- and mannose-capped glycans, and were equally or more efficient than the shedding inhibitors GM6001 and EGTA, we conclude that shear-mediated desialylation of platelet glycans is an upstream signaling event that triggers sequential deglycosylation of underlying glycans and/or enzymatic shedding of now desialylated glycoproteins by metalloproteinases. On other examples, it was also shown that site-specific glycosylation of juxtamembrane regions coregulates metalloproteinase-mediated ectodomain shedding of membrane proteins, affecting their susceptibility to proteolysis. Removal of O-glycans facilitates proteolytic shedding of TNF-α, IL-2 and IL-6 receptors.^36,37^ Similarly, desialylation of O-glycans in GPIb-IX by neuraminidase is associated with abnormal expression on the platelet surface, unfolding of mechanosensitive domain, and extensive shedding of GPIbα ectodomain by metalloproteinase ADAM17.^38^ It is conceivable that, besides GPIb-IX, other highly glycosylated platelet receptors that undergo shear-mediated shedding by metalloproteinases (GPVI, PECAM-1, P-selectin) might first undergo enzymatic deglycosylation and only then be proteolytically cleaved.

Neuraminidases are glycoside hydrolases that cleave glycosidic linkages in Sias and remove Sia caps from targeted glycans. In the human genome, four neuraminidases have been identified (NEU1-4), and all four of them were detected in platelets^35,39^. Yet, very limited information is available regarding subcellular localization of platelet neuraminidases, and their functional role is under intense scrutiny. In resting platelets, NEU1 and NEU3 has been located in granular-like compartments, while NEU2 and NEU4 were detected in some α-granules and on plasma membrane, respectively.^35,39^ Furthermore, early functional studies by Holmsen and colleagues suggest that platelets store and secrete other glycosidases, including galactosidase, N-acetylglucosaminidase, glucuronidase, and acid phosphatase, presumably located in granular compartments other than α- or dense granules.^40,41^ Herein, we demonstrated that platelets possess functionally active extracellular and intracellular glycosidases; some are constitutively active (mannosidase and hexosaminidase), while activity of others is upregulated by shear stress (neuraminidase and galactosidase). We found that resting platelets expressed very high basal activity of *extracellular* mannosidase and hexosaminidase, while neuraminidase and galactosidase activities were barely traceable. Moreover, platelets contained highly active *intracellular* hexosaminidase and galactosidase; low basal activities of intracellular neuraminidase and mannosidase were also detected. Curiously, *extracellular* platelet glycosidases exhibited very high enzymatic activity in neutral and not in acidic pH, which contradicts a prevalent notion that cellular glycosidases reside in lysosomes and have acidic pH optimum. Yet, it is conceivable that platelet surface glycosidases operate within the blood milieu with a strictly regulated pH 7.4^42^, and therefore neutral pH optimum is reasonable. As for *intracellular* glycosidases, galactosidase, hexosaminidase, and mannosidase indeed exhibited high enzymatic activities in acidic pH, suggesting their primarily lysosomal localization in platelets. Nonetheless, intracellular neuraminidase still preferred neutral pH hinting towards alternative subcellular localization. Our findings are in agreement with imaging data by van der Wal et al. showing lack of colocalization of platelet NEU1 and NEU2 with lysosomal marker LAMP1.^39^ Jansen at al. however demonstrated partial colocalization of platelet NEU1 and β-galactosidase, similarly to other cells where α- neuraminidase forms a functional multienzyme complex with β-galactosidase and other lysosomal glycosidases protecting them from lysosomal degradation.^35,43^

Next, we demonstrated that platelet exposure to shear stress resulted in selective upregulation of platelet neuraminidase and galactosidase activities. Enzymatic activities of neutral extracellular neuraminidase and galactosidase post-shear increased 8- and 1.5-times, respectively, as compared to their basal levels. We noticed that shear-mediated upregulation was solely directed towards neutral neuraminidase and galactosidase – the two exoenzymes that aren’t active on resting platelets. At the same time, activities of neutral hexosaminidase and mannosidase that are highly active on resting platelets remained unaffected by shear stress. The selective upregulation of neuraminidase and galactosidase on platelet surface by shear suggests that, unlike other glycosidases, neuraminidase and galactosidase play a unique regulatory role, so their functional activities should be limited (**Fig 6C**). Recently, it has been demonstrated that in refrigerated platelets^35^ and platelets from patients with refractory immune thrombocytopenia^33^, endo-neuraminidases could be translocated and overexpressed on the platelet membrane, promoting subsequent desialylation of platelet glycans and intense proteolytic shedding of GPIb by ADAM17. Upregulation of NEU1 surface expression following refrigeration resulted in the increase of neuraminidase activity on murine platelets that corresponded to 27%-44% of their total neuraminidase activity in permeabilized platelets^35^. Curiously, galactosidase expression and activity also increased on platelet membrane during long-term storage^21,35^. These observations led authors to believe that platelet neuraminidase and galactosidase are both located in so-called secretory lysosomes, a unique subtype of secretory organelles that are finely controlled and released in response to certain stimuli. Our findings tend to agree with this notion. In our hands, exposure to shear stress was associated with a modest increase of P-selectin and tetraspanin CD63 surface expression on platelets, both markers of platelet degranulation and lysosomal release. The minor increase of acidic glycosidase activity on sheared platelets also hints toward shear-mediated release of lysosomal content.

Analyzing alternative sources of glycosidase activity in blood, we found that plasma offers a rich source of active glycosidases, including mannosidase, hexosaminidase, and to a lesser extent neuraminidase. As expected, all plasma glycosidases demonstrated high enzymatic activity in neutral pH. Hexosaminidase was also equally active in acidic pH suggesting that two separate iso-enzymes with different pH optimums are present in plasma. In PRP, mannosidase and hexosaminidase activities were elevated due to complementary contribution from plasma- and platelet-bound pools of these glycosidases. The same cannot be said about neuraminidase and galactosidase. Since neuraminidase activity was only present in plasma and not on unstimulated platelets, no increase of neuraminidase activity was observed in PRP as compared to PPP. In lysed PRP however, both neutral and acidic neuraminidase activities increased slightly due to release of an intraplatelet neuraminidase pool. Curiously, no galactosidase activity was registered in plasma. Nonetheless, neutral galactosidase activity appeared in PRP at levels comparable to intact GFP, indicating that in healthy individuals, platelets are primary carriers of galactosidase activity. Furthermore, in lysed PRP, strikingly high acidic galactosidase activity was detected, likely originating from intracellular platelet storages. Our findings tend to line up with prior reports. Analyzing glycosidase activities in healthy and septic individuals at *neutral* pH, Haslund-Gourley et al. showed comparably high mannosidase and hexosaminidase activities in plasma of healthy individuals, while galactosidase activity was rather low.^42^ Neutral hexosaminidase and galactosidase activities (but not mannosidase) significantly increased in septic patients, with elevated galactosidase activity showing prognostic indications for the increased mortality rate. Very high *acidic* hexosaminidase activity and much lower *acidic* mannosidase and galactosidase activities were previously detected in human plasma.^29^ Due to different activity units used in the two studies, it is difficult to compare and evaluate the contribution of neutral versus acidic glycosidase pools. One report has shown that, in murine plasma and serum, acidic glycosidase activity was significantly higher than neutral activity for galactosidase, hexosaminidase, and mannosidase.^42^ While this finding contradicts our observations, we suspect that intra-species differences in plasma enzymes’ content and kinetics could be to blame for such a discrepancy.

We evaluated the glycosidase activity in serum obtained from whole blood where clotting was facilitated by recalcification. We hypothesized that glycosidase activities in serum should be higher than in plasma due to potential release of platelet-bound glycosidases following platelet activation by thrombin generated during coagulation. We demonstrated that neuraminidase and galactosidase activities in serum was indeed significantly higher than in plasma. Interestingly, only neutral neuraminidase activity increased in serum, while for serum galactosidase both neutral and acidic activities were elevated. Similarly, enzymatic activities of acidic (but not neutral) hexosaminidase and mannosidase were nearly twice as high in serum as compared to plasma activity levels. Previous reports also recorded increased glycosidase activity in serum as compared to plasma in humans and mice, and the extent of the increase drastically varied for different glycosidases^29,42^ and for acetic and neutral isoenzymes of the same glycosidase^42^. Next, we noticed that for neutral neuraminidase, neutral galactosidase, and acetic mannosidase the incremental increase of their activity in serum (as compared to plasma) was equivalent to platelet activity of these enzymes, while for acetic galactosidase platelets exhibited much higher intracellular acetic galactosidase activity than was detected in serum. Furthermore, following clotting and shear exposure, neuraminidase and galactosidase are released from platelets to a greater extent than mannosidase and hexosaminidase. Holmsen & Weiss have previously demonstrated that up to 30-60% of platelet acidic glycosidases are released when platelets are stimulated with strong agonists, including thrombin and collagen, and the release patterns of various glycosidases differ from one another.^41^ Cumulatively, these observations strongly suggest that upon activation platelets selectively release their highly active glycosidases in blood milieu, potentially affecting glycosylation status of blood cells and proteins, and various secretory mechanisms are operative for various platelet glycosidases and/or their isoenzymes.

Lastly, we studied the effect of shear stress and neuraminidase on platelet count and microvesiculation. Exposure to shear stress of increasing magnitude and duration rendered an incremental decrease in platelet count and associated increase in PDMPs, as indicated by two platelet-specific markers CD41 (αIIbβ3) and CD42a (GPIX). Exogenous neuraminidase further exacerbated shear-mediated decrease in platelet count and facilitated shear-mediated microvesiculation leading to up 30% and 50% increase in PDMPs following 30 and 70 dyne/cm^2^ shear, respectively. Interestingly, synergistic effects of neuraminidase on shear-mediated platelet decline and increased microvesiculation were more pronounced for higher shear magnitude. We also noticed that shear stress induced a minor decrease in platelet αIIbβ3 surface density and had no significant effect or even slightly increased platelet GPIX. Nonetheless, sheared PDMPs exhibited significantly higher surface densities of both αIIbβ3 (2.4x) and GPIX (3.4x) than platelets, thus suggesting that these receptors were redistributed from platelet surface to microparticles during shear-mediated microvesiculation. This observation lines up with our finding that PDMPs demonstrated significantly higher surface glycosylation levels than platelets and further supports the idea that glycoprotein receptors, such as αIIbβ3 and GPIb-IX, are primary carriers of platelet surface glycosylation. Unlike shear, neuraminidase induced significant upregulation of αIIbβ3 and GPIX surface expression on platelets and microparticles. Neuraminidase- induced increase of αIIbβ3 expression was resistant to shear stress and did not decrease even after 70 dyne/cm^2^ shear exposure. GPIX increased consistently 50-69% of its surface density on unsheared and sheared platelets alike after neuraminidase treatment. Neuraminidase-mediated upregulation of αIIbβ3 and GPIX coincided with the intensification of platelet granule release as indicated by the increase of P-selectin and tetraspanin CD63 on platelets following exposure to neuraminidase and shear stress, but not neuraminidase alone. Similarly, Kullaya et al. demonstrated that platelet exposure to pneumococcal NeuA and thrombin, and not NeuA alone, resulted in elevated surface expression of platelet β3 (CD61) and secretion of dense granules as indicated by upregulation of CD63 on platelets and decrease of intraplatelet mepacrine concentration.^31^ NeuA also promoted agonist-induced fibrinogen binding and P-selectin surface expression, leading authors to conclude that platelet desialylation was associated with platelet hyperactivity. While increased platelet degranulation could explain the hyperactivity of NeuA-desialylated platelets and intensification of microvesiculation following neuraminidase treatment observed in our study, it can’t explain intensification of platelet count drop when neuraminidase-treated platelets were exposed to shear stress. We and others have previously demonstrated that shear-stress induces proapoptotic platelet behavior, associated with mitochondrial collapse, caspase activation, shedding of platelet adhesion receptors, and intense microvesiculation.^12,14,15^ Concurrently, it has been documented that intense platelet deglycosylation and shedding of glycoprotein receptors is associated with intensification of platelet apoptosis and rapid clearance from the circulation under a number of pathological conditions, including immune thrombocytopenia, sepsis, viral and microbial infections.^31–33^ Yet, direct link between platelets desialylation and apoptosis hasn’t been established. Recent studies have shown that NEU1 plays a crucial role in regulation of mitochondrial metabolism in cardiomyocytes, and upregulation of NEU1 and β- galactosidase expression is characteristic for post-infarction cardiomyocytes and infiltrating monocytes and macrophages after ischemia-reperfusion.^44,45^ The cardiomyocyte specific NEU1 knockdown alleviated mitochondrial energy metabolism disorder, oxidative stress, and myocardial remodeling induced by infarction.^44^ Taking in consideration these data, we do not exclude the possibility that shear-mediated upregulation of neuraminidase activity in platelets could facilitate mitochondrial collapse and further induce platelet apoptosis. Yet, to verify this hypothesis further research is required.

Regarding limitations, routine flow cytometry used to capture and characterize sheared PDMPs underestimates smaller microparticle (<300 nm) and exosome (<100 nm) populations,^46^ as such these particles were under-accounted for in our study. It’s reasonable to assume that lectin blotting underestimates the prevalence of glycosylated proteins in platelets, only revealing the most highly expressed and highly glycosylated ones. LCA lectin used to detect terminal mannose can also bind terminal fucose.^27^ Therefore, mannose levels reported herein might be in part attributed to fucose.

## AFFILIATIONS

Departments of Medicine (Y.R.-M., S.L., E.C.) and Biomedical Engineering (M.J.S.), Sarver Heart Center, University of Arizona, Tucson. Arizona Center for Accelerated Biomedical Innovation, University of Arizona (Y.R.-M., M.J.S.). Vascular Biology Program, Boston Children’s Hospital, Harvard Medical School, Boston, MA (J.E.I.).

## ACKNOWLEDGMENTS

1. Y. Roka-Moiia designed the study, performed experiments, analyzed and interpreted data, wrote and edited the article, and acquired funding; S. Lewis and E. Cleveland performed experiments; J.E. Italiano participated in discussions and edited the article; M.J. Slepian participated in discussions, edited the article, and acquired funding.

## NONSTANDARD ABBREVIATIONS AND ACRONYMS

DANA: 2-deoxy-N-acetylneuraminic acid
ECA: *Erythrina Cristagalli* agglutinin
ECMO: extracorporeal membrane oxygenation
GalNAc: *N-*acetylgalactosamine
GFP: gel-filtered platelets
GlcNAc: *N-*acetylglucosamine
GSL: *Griffonia Simplicifolia* lectin II
LCA: *Lens Culinaris* agglutinin
MAL: *Maackia Amurensis* lectin II
MCS: mechanical circulatory support
PDMPs: platelet-derived microparticles
PPP: platelet-poor plasma
PRP: platelet rich-plasma
RCA: *Ricinus Communis* agglutinin I
SBA: soybean agglutinin
Sias: sialic acids
SMPD: shear-mediated platelet dysfunction
SNA: *Sambucus Nigra* agglutinin I
VAD: ventricular assist device

